# Transcriptomics and weighted protein network analysis of the LRRK2 protein interactome reveal distinct molecular signatures for sporadic and LRRK2 Parkinson’s Disease

**DOI:** 10.1101/2023.09.12.557373

**Authors:** Yibo Zhao, Matthew Bracher-Smith, Kirsten Harvey, Valentina Escott-Price, Patrick A Lewis, Claudia Manzoni

## Abstract

Mutations in the LRRK2 gene are the most common genetic cause for familial Parkinson’s Disease (LRRK2-PD) and an important risk factor for sporadic PD (sPD). Multiple clinical trials are ongoing to evaluate the benefits associated with the therapeutical reduction of LRRK2 kinase activity. In this study, we described the changes on transcriptomic profiles (whole blood mRNA levels) of LRRK2 protein interactors in the sPD and LRRK2-PD cases as compared to healthy controls with the aim of comparing the two PD conditions. We went on to model the protein-protein interaction (PPI) network around LRRK2, which was weighted to reflect the transcriptomic changes in the network based on the expression and co-expression levels. Our results showed that LRRK2 interactors present both similar but also different alterations in expression levels and co-expression behaviours in the sPD and LRRK2-PD cases. The similar changes result in decreased connectivity within a topological cluster of the LRRK2 PPI network associated with ribosomal functions and DNA/RNA metabolism in both the sPD and LRRK2-PD scenario; while the connectivity within the autophagy/mitophagy/neurotransmitter-transport related cluster was increased exclusively in the LRRK2-PD condition as compared to the healthy controls. These results suggest that, albeit being classified as the same disease based on clinical features, LRRK2-PD and sPD show some significant differences from a molecular perspective.

## Introduction

Leucine-rich repeat kinase 2 is a large (over 250 kDa), multifunctional enzyme encoded by the LRRK2 gene, possessing 2 enzymatic (GTPase and Kinase) and 4 scaffold (Armadillo, Ankyrin, LRR and WD40 motifs) domains (Berwick et al., 2019). LRRK2 is able to interact with a vast number of protein partners (Zhao et al., 2023) and has been implied in a large number of biological processes, including vesicular transport, autophagy, regulation of cellular response to stress, regulation of cell cycle, etc (Albanese et al., 2019; Chen et al., 2017; Hsu et al., 2010; Sanna et al., 2012). Mutations in LRRK2 are an important genetic cause of familial PD (fPD) overall with 1 to 40% of fPD cases associated with LRRK2 depending on the population under study (Bras et al., 2005; Greggio et al., 2008; Lesage et al., 2006; Ozelius et al., 2006). Since 2004, when a missense change on the LRRK2 gene was firstly associated with fPD, numerous coding and non-coding variants of LRRK2 have been identified in PD families, among which the G2019S and R1441C/G mutations are the 2 most common pathogenic variants occurring on the kinase and GTPase domains of the LRRK2 protein respectively (Dauer & Ho, 2010; Esteves et al., 2014; Giesert et al., 2017; Henry et al., 2015; Paisán-Ruiz et al., 2013). However, how these protein coding changes contribute to the aetiology of fPD is still unclear. Additionally, non-coding polymorphisms, mainly in the promoter of LRRK2, have been linked to sporadic PD (sPD) (Nalls et al., 2019) and upregulated LRRK2 kinase activity has also been related with sPD. For example, PD-related inflammation both in the Central Nervous System (CNS) and at the periphery, was linked to an increased LRRK2 expression level and strengthened LRRK2 kinase activity in microglia and peripheral immune cells in sPD patients as compared to controls (Cook et al., 2017; Di Maio et al., 2018). All the above-mentioned findings indicate that LRRK2 is crucial for the understanding of PD etiopathogenesis and supports the thesis that LRRK2 might constitute a link between familial and sporadic forms of the disease.

However, from a clinical perspective, LRRK2-PD and sPD have been reported with some distinct features. Despite having similar motor-symptoms (such as bradykinesia, tremor, rigidity, and postural instability) as well as sharing some of the non-motor symptoms (Haugarvoll et al., 2008; Healy et al., 2008; Kluss et al., 2019), patients with LRRK2-PD show slower decline considering both movement and cognitive impairment (Alcalay et al., 2015; Srivatsal et al., 2015). In addition, LRRK2-PD and sPD present with distinct pathological features. For example, LRRK2-PD patients exhibit less α-synuclein aggregation in Cerebrospinal Fluid (CSF), feature that is a hallmark of sPD (Garrido et al., 2019; Rivero-Ríos et al., 2020) as well as increased basal forebrain volume, which may represent a compensation in the cholinergic system (Batzu et al., 2023). Such differences might highlight an intrinsic variation at the molecular level between these two forms of PD thus suggesting different model systems might be required to investigate them. Also, this consideration may pose a problem in translational research, for example the use of LRRK2 inhibitors in clinical trials (Tolosa et al., 2020) might require patient stratification.

In this study we hypothesized that despite being generally regarded as the same disease, sPD and LRRK2-PD might have a different molecular signature and therefore the molecular alterations contributing to disease onset and progression might be functionally different. We constructed a protein-protein interaction (PPI) network around LRRK2 (LRRK2_net_) and evaluated the mRNA level changes within the LRRK2_net_ in a cohort of sPD and LRRK2-PD patients in comparison with healthy controls. The results provide bioinformatic evidence that, despite some similarities, the signature of expression changes in the LRRK2_net_ is overall diffferent in sPD vs LRRK2-PD showing significant differences in both gene expression and co-expression, suggesting the molecular pathways at the base of these two conditions might be different. This is relevant for the understanding of the different molecular mechanisms of PD, and it highlights the necessity for patient stratification in both discovery research and clinical trials, suggesting different therapeutic approaches might be needed if we intend to move from symptomatic to effective disease treatment.

## Method

### LRRK2 protein interactors download and quality control (QC)

LRRK2 protein interactors were downloaded via PINOT v1.1 (http://www.reading.ac.uk/bioinf/PINOT/PINOT_form.html), HIPPIE v2.3 (http://cbdm-01.zdv.uni-mainz.de/~mschaefer/hippie/index.php) and MIST v5.0 (https://fgrtools.hms.harvard.edu/MIST/) (Alanis-Lobato et al., 2017; Hu et al., 2018; Tomkins et al., 2020) on 16^th^ March 2023 and the LRRK2 interactome was built following the pipeline in (Zhao et al., 2023). To access the most comprehensive set of LRRK2 interactors, “Lenient” filter level was applied in PINOT; while no filter was applied for HIPPIE and MIST. Interactors retrieved from the 3 tools were merged and QC-ed to identify interactors with missing publication identifier, missing interaction detection method, no conversion to a standard gene identifier, and with low interaction confidence score.

### Whole blood RNA-Seq data download and QC

Baseline (BL = time at diagnosis) whole blood mRNA data (read counts) of LRRK2 interactors for healthy controls (HC), sPD patients and LRRK2-PD patients were retrieved using Ensembl gene ID from the Parkinson’s Progression Marker Initiative (PPMI) dataset on 24^th^ March 2023. PPMI is an ongoing observational, international, multicentred cohort study aimed at identifying the biomarkers of PD progression in a large cohort of participants (https://www.ppmi-info.org/). It is a public-private partnership – is funded by The Michael J. Fox Foundation for Parkinson’s Research and funding partners, including those reported at https://www.ppmi-info.org/about-ppmi/who-we-are/studysponsors. Subject QC: PPMI cohorts of “de novo PD” and “healthy control” were included in this study. Subjects from the 2 cohorts were further filtered to keep only those with robust genetic status records using the following criteria: confirmed by at least 3 out of 6 detection techniques (WGS, WES, RNA-Seq, GWAS, CLIA, SANGER) of which 1 should be a next generation sequencing technique (WGS, WES, RNA-Seq) and 1 should be a screening technique (GWAS, CLIA, SANGER). Of note, subjects with mutations in PD genes other than LRRK2 (GBA, PINK, Parkin and SNCA) were excluded to avoid potential bias. QC-ed subjects were allocated into 3 cohorts: Control, Sporadic PD (sPD, with no LRRK2 or other gene mutations in record) and LRRK2-PD (only with LRRK2 mutations considered to be pathogenetic – i.e., G2019S, R1441C/G). In addition, Principal Component Analysis (PCA) was performed to remove potential outlier subjects. Transcript QC: Transcripts of LRRK2 interactors with read counts ≤ 15 in more than 75% samples were removed (Langfelder & Horvath, 2008). Read counts of LRRK2 interactors retrieved from PPMI were extracted for the 3 cohorts, thereby forming the “PPMI_Matrix”.

### Differential Expression Analysis and classification models for sPD and LRRK2-PD

The PPMI_Matrix was then normalised via the median of ratios method using the “count” function in the R package “DESeq2” (Love et al., 2014). The normalised PPMI_Matrix (hereby referred as “norm_PPMI_Matrix”) was utilised to perform Differential Expression Analysis (DEA) to compare the expression levels of LRRK2 interactors in the control, sPD and LRRK2-PD cohorts using “DESeq2”. Fold change (FC) for each of the LRRK2 interactors (*i*) in [sPD vs control] and (*ii*) in [LRRK2-PD vs control] was calculated. P-value adjustment for multiple comparisons were performed via Bonferroni’s method. The results from DEA were further adjusted for sex. LRRK2 interactors were considered up(down)-regulated when log2FC > 0 (log2FC < 0) and adjusted-p < 0.05 in [sPD vs control] or [LRRK2-PD vs control]. Up/down-regulated LRRK2 interactors in the 2 PD conditions were functionally annotated via Gene Ontology Biological Process (GO-BP) enrichment analysis. The read counts of LRRK2 interactors with significant up/down-regulated expression in the 2 PD conditions as compared to controls were utilised to construct a machine learning model via Least Absolute Shrinkage and Selection Operator (LASSO) algorithm via the R package “glmnet”. In order to reduce the risk for model overfitting, univariate logistic regression was performed on each LRRK2 interactor and only those with p-value < 0.05 were included in the LASSO regression model. The train-test split ratio for the LASSO regression model was set as 4:1. The tunning parameter lambda (λ) were optimised by a 10-fold cross-validation (CV) to reach the minimum Mean-Squared Error (MSE) via the “cv.glmnet” function of the “glmnet” package. The refined models were then assessed on the test set. Receiver Operating Characteristic (ROC) curves were generated via the “roc.glmnet” function of the “glmnet” package.

### Weighted Gene Co-expression Network Analysis

Signed Weighted Gene Co-expression Network Analyses (WGCNA) were performed on the norm_PPMI_Matrix via the R package “WGCNA” to identify co-expression modules within the LRRK2_net_ across the sPD, LRRK2-PD and control conditions. Module-Trait correlation was evaluated via the “corPvalueStudent” function in the “WGCNA” package.

### LRRK2 PPI network (LRRK2**_net_**) construction and weighted network analysis

To construct the LRRK2_net_, the 2^nd^-layer PPIs (i.e., PPIs among LRRK2 interactors) were downloaded via HIPPIE (v2.3) on 16^th^ March 2023. The 2^nd^-layer PPIs with high confidence score (≥ 0.72) were kept for network construction (the LRRK2_net_). The Fast Greedy Clustering algorithm (Newman, 2004) was utilised to detect topological clusters in the LRRK2_net_ based on edge betweenness (i.e., calculating the number of shortest paths between any pair of nodes in the network that pass-through a given edge), via the “cluastermaker2” Cytoscape add-in (v2.3.4). For each obtained topological cluster, edges were classified as up-, down-regulated or unchanged based on the following criteria: A) up-regulated edge: i) at least 1 of the 2 nodes connected by the edge had increased expression level in sPD and/or LRRK2-PD vs controls or ii) a strong positive co-expression (Pearson’s coefficient > 0.6) was observed for the 2 nodes connected via the edge in sPD and/or LRRK2-PD but not in controls; B) downregulated edge: i) at least 1 of the 2 nodes connected by the edge presented decreased expression level in sPD and/or LRRK2-PD vs controls or ii) a strong positive co-expression (Pearson’s coefficient > 0.6) for the 2 nodes connected via the edge was observed in the controls but not in sPD and/or LRRK2-PD cases. The percentage of upregulated, downregulated and unchanged edges for each single topological cluster were calculated for the sPD and the LRRK2-PD scenarios and compared via One Sample Proportion Test to identify the trend of each topological cluster and qualitatively define whether a cluster was mainly up/down regulated or unchanged in sPD or LRRK2-PD vs controls.

### Functional enrichment analysis

In this study, functional enrichment analyses for LRRK2 interactors were performed via the webtool “g:Profiler” (https://biit.cs.ut.ee/gprofiler/gost) (Peterson et al., 2020).The parameters were set as following: organism - Homo sapiens (Human); data source - GO biological process (GO-BPs) only; statistical domain scope – annotated genes only; statistical method - Fisher’s one-tailed test; significance threshold – Bonferroni correction (threshold = 0.05). No hierarchical filtering was included. To increase the sensitivity of analysis, a cut-off of ≤ 2500 was set for the “term size” of enriched GO terms.

## Results

### Whole-blood transcriptomic profiling for LRRK2 interactors

A total of 418 protein interactors of LRRK2 were retrieved via an in-house pipeline (Zhao et al., 2023) (**Table S1**). Whole blood RNA-Seq read counts were retrieved for 415 (out of 418) LRRK2 interactors and for 657 PPMI subjects with validated genotyping data (controls = 170; sPD cases = 371; and LRRK2-PD cases = 116). Demographic features of the 2 included cohorts were listed in Table 1. A total of 38 interactors were removed due to low read counts, and no subjects were identified as outliers by PCA (**Figure S1**). Hence, mRNA levels of the remaining 377 LRRK2 interactors of 657 PPMI subjects formed the PPMI_Matrix. DEA results showed that: the mRNA levels of 64 interactors (17.0%) were significantly altered in the LRRK2-PD cases vs. controls, with 36 down-regulated and 28 up-regulated interactors (adjusted-p < 0.05, **Figure 1A**, **Table S2**). A total of 53 interactors (14.0%) presented significant changes in expression levels in sPD in comparison with controls, including 29 down-regulated and 24 up-regulated interactors (adjusted-p < 0.05, **Figure 1B**). Of note, only a total of 17 (5.6%) interactors presented with the same alteration in LRRK2-PD and sPD, in which 14 interactors were down-regulated while 3 interactors were up-regulated, suggesting these LRRK2 interactors were consistently affected during PD progression regardless of the existence of a *LRRK2* mutation (**Figure 1C**). Functional enrichment analysis associated these 17 proteins with GO-BP enrichment analysis showed that these 17 interactors were associated with biosynthesis, and protein metabolism (**Figure 1D**). LRRK2 interactors which were up-regulated in the LRRK2-PD cases were enriched for protein and DNA metabolism, synaptic transmission, cell death, response to stress and response to misfolded protein, while the ones up-regulated in the sPD cases contributed to protein metabolism, signalling, intracellular organisation, response to stress, cell cycle and autophagy (**Table S3**). In comparison, LRRK2 interactors which were down-regulated in the LRRK2-PD cases were related to biosynthesis, while those down-regulated in the sPD cases were associated with protein metabolism, cell death and biosynthesis (**Table S3**). These findings suggest that the sPD and LRRK2-PD pathology may affect different sets of LRRK2-mediated biological processes.

**Table 1.** PPMI cohort characterisation

|  | **Control** | **sPD** | **LRRK2-PD** |
| --- | --- | --- | --- |
| Gender (Male) | 94 (55.3%) | 224 (60.4%) | 71 (61.2%) |
| Age (Mean (SD)) | 60.9 (0.8) | 61.3 (0.5) | 63.1 (0.8) |
| Ethnicity (White) | 160 (94.1%) | 360 (97.0%) | 108 (93.1%) |
| MDS-UPDRS III | - | 21.1 (0.5) | 20.1 (0.9) |

*Note: Movement disability for the sPD and LRRK2-PD cases was evaluated via MDS-UPDRS III (Score range: 0–132; 32 and below is mild, 59 and above is severe) (Balestrino et al., 2019; Skorvanek et al., 2017). Abbreviations: MDS-UPDRS: Unified Parkinson's Disease Rating Scale published by the Movement Disorder Society*

**Figure 1.**
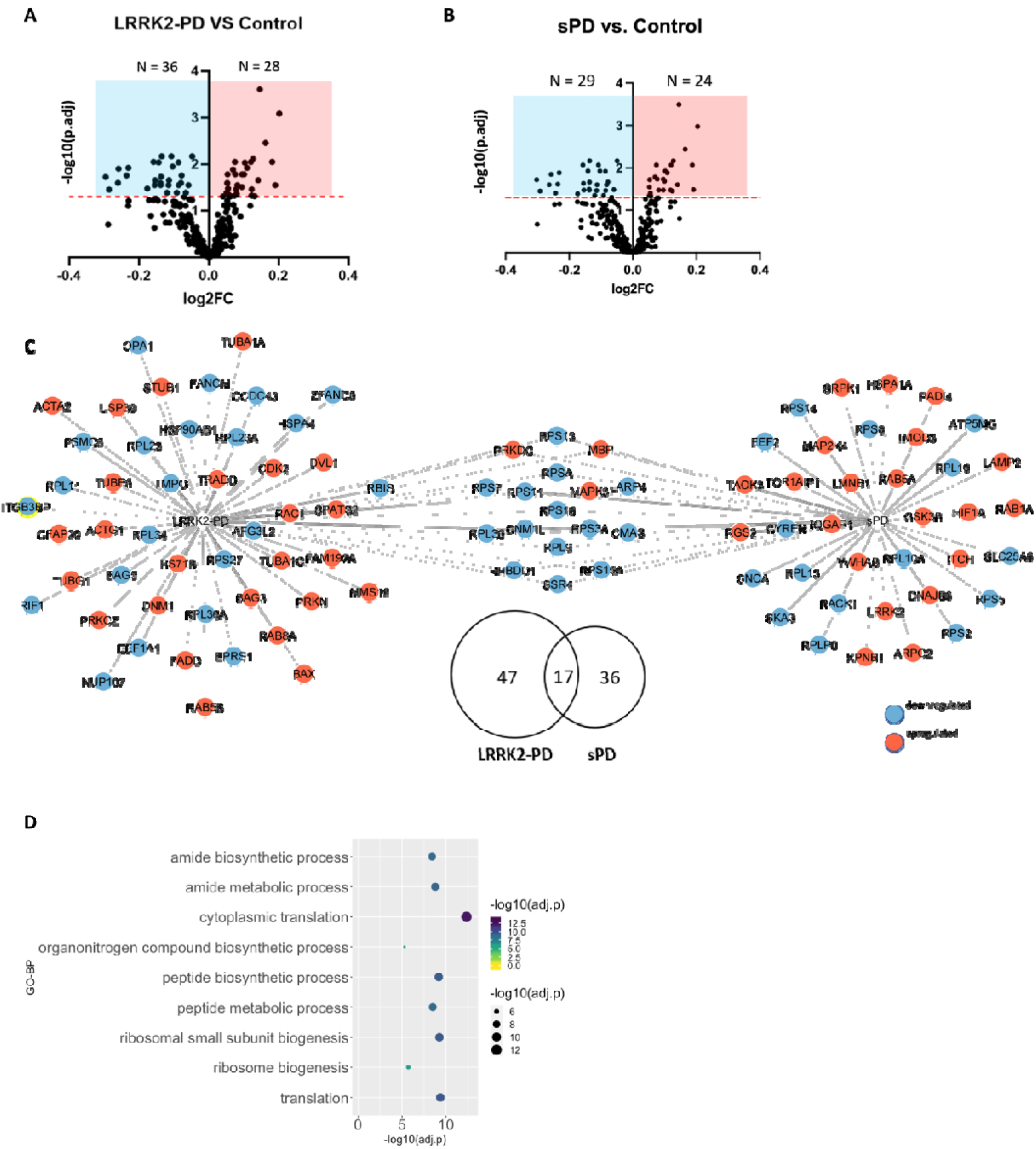
DEA on LRRK2 interactors in the LRRK2-PD and sPD cases vs. Controls. **A)** The volcano plot shows DEA results for the 377 LRRK2 interactors with available whole blood RNA expression levels in the PPMI dataset comparing the LRRK2-PD cohort vs controls. The X-axis represents log2 Fold Change (log2FC); the Y-axis represents -log10 transformed adjusted-p values (-log10(adj.p)). The red horizonal line represents the threshold of adjusted-p = 0.05. LRRK2 interactors in the red area (on the right side) were considered up-regulated (N = 28) while those in blue area (on the left side) were considered down-regulated (N = 36). **B)** The volcano plot shows DEA results for LRRK2 interactors comparing the sPD cohort vs controls. LRRK2 interactors in the red area (on the right side) were considered up-regulated (N = 24) while those in blue area (on the left side) were considered down-regulated (N = 29). **C)** The Venn graph and the network graph show 17 LRRK2 interactors presenting same differential expression pattern in the LRRK2-PD and the sPD cohorts in comparison with controls. In the network graph, interactors with significant differential expression profiles were colour-coded based on up-regulation (red) and down-regulation (blue). **D**) The bubble graph shows the GO-BP terms enriched for the 17 LRRK2 interactors that presented similar alterations in the 2 PD conditions as compared to the Controls. Bubble colour and size refer to the enrichment significance (evaluated as -log10 transformed adjusted p-values, - log10(adj.p)).

### Transcriptomic features of LRRK2 interactors differentiated the LRRK2-PD and sPD cases

Univariate logistic regression was performed on each of the 100 LRRK2 interactors with significant differential expression in the sPD and/or LRRK2-PD cohorts vs. controls, out of which a total of 14 interactors with p-value < 0.05 were selected for further model construction, including STUB1, DVL1, ACTA2, CDK2, MMS19, PRKN, TUBB6, TUBG1, BAG3, HSPA1A, LMNB1, SNCA, RPS2 and SLC25A6 (**Table 2**). A LASSO regression model constructed with the mRNA levels of the 14 LRRK2 interactors was then trained on a randomly-picked cohort of 296 sPD cases and 92 LRRK2-PD cases. A λ value of 0.008 (log(λ) = -4.869 was chosen to reach the minimum MSE = 0.171 (Figure 2A), leaving a total of 8 interactors in the model, including STUB1 (coefficient = -1.61), ACTA2 (coefficient = -0.85), PRKN (coefficient = = -0.38), TUBB6 (coefficient = -1.21), HSPA1A (coefficient = 3.12), LMNB1 (coefficient = 0.55), SNCA (coefficient = -0.46) and SLC25A6 (coefficient = -0.68) (Figure 2B). The cut-off value on the predicted value was optimised as 0.94 to reach the maximum accuracy in the training set (AC_train = 72.9%, Figure 2C). The refined model was then validated on the test set, containing 75 sPD cases and 24 LRRK2-PD cases. ROC curve showed an AUC = 0.68 (95% CI: 0.56-0.80), suggesting a good classification performance of the LASSO model built upon the mRNA levels of LRRK2 interactors (Figure 2D).

**Table 2.** LRRK2 interactors selected by univariate analyses for LASSO model.

| Interactor | Odd Ratio (OR) | 95% Confidence Interval (CI) |  | p-value |
| --- | --- | --- | --- | --- |
|  |  | Upper Limit | Lower Limit |  |
| STUB1 | 0.24 | 0.06 | 0.91 | 0.036 |
| DVL1 | 0.17 | 0.04 | 0.68 | 0.012 |
| ACTA2 | 0.31 | 0.13 | 0.77 | 0.012 |
| CDK2 | 0.22 | 0.06 | 0.86 | 0.029 |
| MMS19 | 0.23 | 0.06 | 0.83 | 0.025 |
| PRKN | 0.18 | 0.06 | 0.56 | 0.003 |
| TUBB6 | 0.18 | 0.06 | 0.49 | 0.001 |
| TUBG1 | 0.25 | 0.07 | 0.90 | 0.035 |
| BAG3 | 0.18 | 0.05 | 0.61 | 0.006 |
| HSPA1A | 3.23 | 1.25 | 8.34 | 0.016 |
| LMNB1 | 2.92 | 1.14 | 7.44 | 0.025 |
| SNCA | 0.45 | 0.21 | 0.96 | 0.040 |
| RPS2 | 0.29 | 0.10 | 0.86 | 0.026 |
| SLC25A6 | 0.24 | 0.07 | 0.84 | 0.025 |

**Figure 2.**
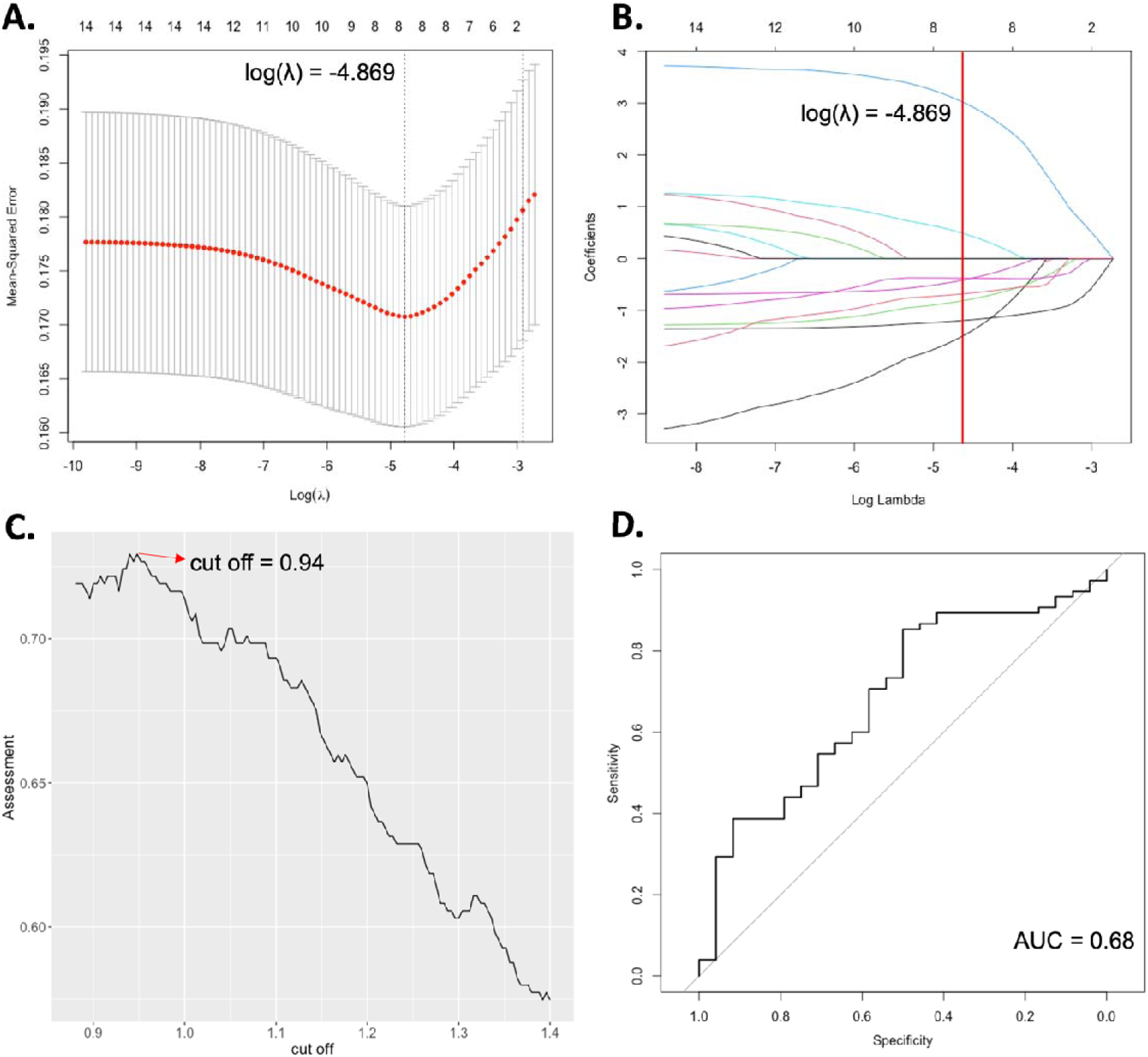
Classification model for the sPD/LRRK2-PD differentiation based on the transcriptomic features of LRRK2 interactors. **A)** The Logistic regression model with LASSO (Least Absolute Shrinkage and Selection Operator) was adopted to further reduce the dimensionalities and to select the most significant expression profiles of LRRK2 interactors to differentiate sPD and LRRK2-PD. A λ value of 0.008, with log(λ) = -4.869 was selected according to 10-fold cross-validation; **B)** LASSO coefficient profiles of 14 LRRK2 interactors are plotted. The optimal coefficient profile was produced against the selected λ (marked as the vertical red line), comprised of 8 LRRK2 interactors, namely STUB1 (coefficient = -1.61), ACTA2 (coefficient = -0.85), PRKN (coefficient = = -0.38), TUBB6 (coefficient = -1.21), HSPA1A (coefficient = 3.12), LMNB1 (coefficient = 0.55), SNCA (coefficient = -0.46) and SLC25A6 (coefficient = -0.68). **C)** The distribution curve shows different cut-off values and the model performance (as assessed by the accuracy) on the train set. A cut-off of 0.94 was selected to reach the highest accuracy of 72.9%. **D)** The graph shows the ROC curve of the model validation on the test set, which possessed an AUC value of 0.68.

### Co-expression modules of LRRK2 interactors in the sPD and LRRK2-PD conditions

A soft power β = 28 was selected via the R package “WGCNA” to construct a scale-free signed gene co-expression network (Figure 3A), within which a total of 2 co-expression modules were identified across the 3 examined cohorts: MTurquoise (N = 33 interactors), MBlue (N = 27 interactors) (Figure 3B, **Table S4**). Of note, LRRK2 was found in neither of these 2 modules, suggesting that the overall co-expression level between LRRK2 and its interactors was relatively low in the whole blood, which is in accordance with the findings in our previous study (Zhao et al., 2023). Functional enrichment analysis showed that MTurquoise was related with protein biosynthesis, ribosomal functions and negative regulation of protein ubiquitination; while MBlue was enriched for intracellular organisation, protein phosphorylation and vesicular transport (**Table S5**). Module-Trait correlation analysis showed that the eigengene of MTurqouise (METurqouise) was significantly down-regulated in the LRRK2-PD and sPD cases as compared to control cohort (p < 0.05), while no significant changes were identified for the MEBlue in the 2 PD conditions vs. controls (Figure 3C).

**Figure 3.**
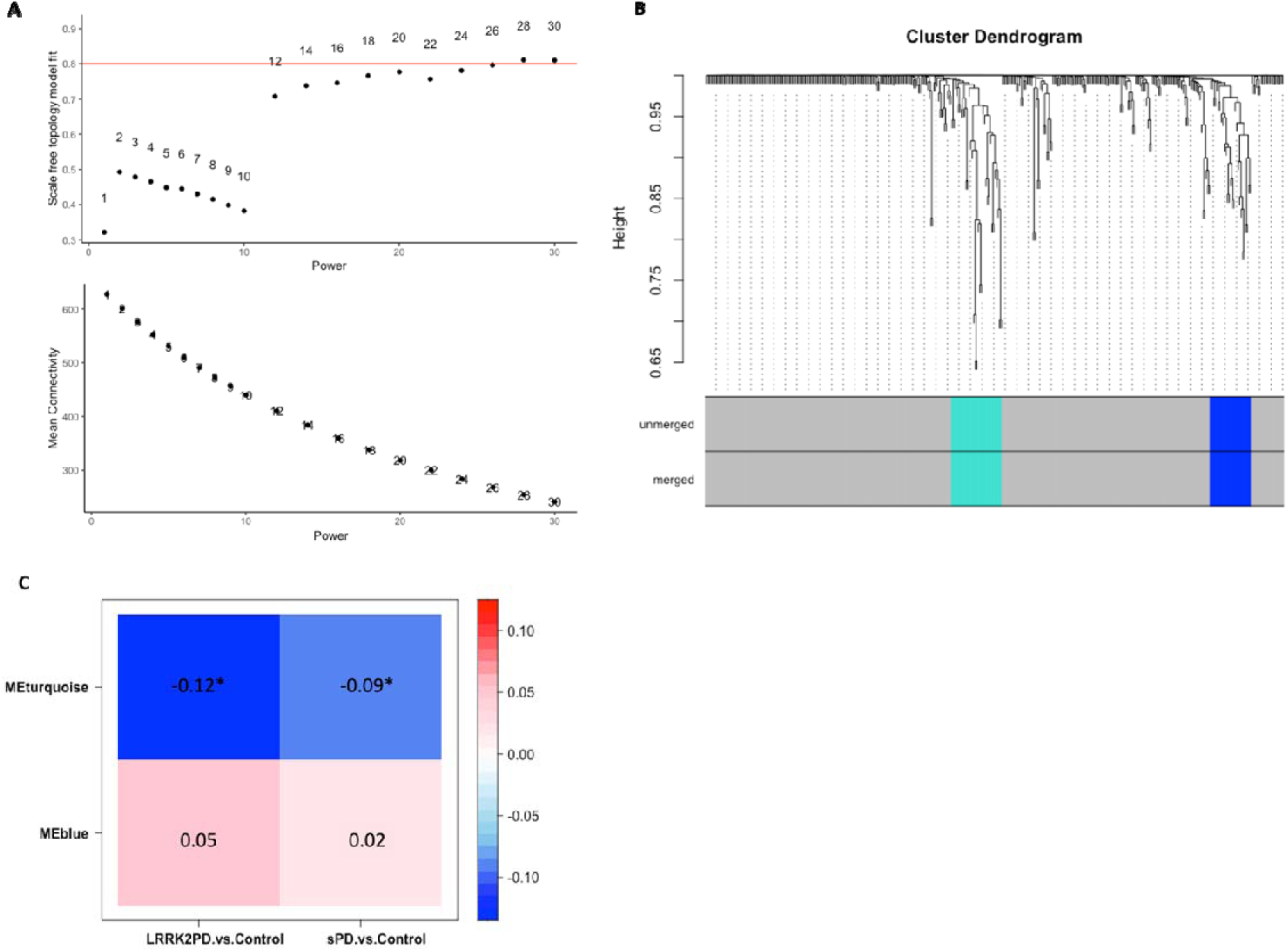
WGCNA on LRRK2 interactors in the sPD, LRRK2-PD and control conditions. **A)** The scatter plots show the selection of a soft power β = 28 was selected for the signed co-expression network constructed among LRRK2 interactors across the 3 cohorts, which achieved scale free model fit and a low mean connectivity. **B)** The dendrogram shows the 2 co-expression modules identified among LRRK2 interactors across the 3 cohorts. Modules are represented by colours (MTurquoise and MBlue). **C)** The heatmap shows the Module-Trait correlation between the eigengene of the 2 co-expression modules (MEturquoise and MEblue) and PD type. The numbers in cells and cell colours represent Pearson’s coefficient. Significant correlation was defined as Pearson’s p-value < 0.05 (marked with *).

### Construction of the LRRK2**_net_**

A total of 4860 “2^nd^-layer” PPIs were extracted from the HIPPIE database (v2.3), among which 1466 (30.2%) were scored as “high confidence” (HIPPIE confidence score ≥ 0.72), out of which 121 self-interactions were removed from the list, thereby leaving 1345 “2^nd^-layer” PPIs for 338 LRRK2 interactors for LRRK2_net_ construction (Figure 4A, **Table S6**). Degree (i.e., the number of PPIs connected to a given interactor) distribution analysis showed that 216 interactors (51.7%) had degrees ≤ 4; 141 interactors (33.7%) presented degrees between 5 and 14; N = 41 interactors (9.8%) presented degrees between 15 and 24; N = 24 interactors with degree ≥ 24, suggesting that the LRRK2_net_ follows the Power Law distribution (log-log plot R-square = 0.8606) (Figure 4B-C, **Table S7**). Interactors with degree ≥ 24 (the top 5% of all) were defined as “sub seed” proteins in the LRRK2_net_, with TP53 (degree = 68), CDK2 (degree = 48), HSPA8 (degree = 46), HSP90AB1 (degree = 44), HSP90AA1 (degree = 43), YWHAZ (degree = 43), LAPR7 (degree = 39), NPM1 (degree = 37), TRAF2 (degree = 32), IQGAP1 (degree = 32), LIMA1 (degree =31), CAPZA2 (degree = 31), PRKN (degree = 28), DBN1 (degree = 28), YWHAQ (degree = 27), RPS8 (degree = 27), YWHAG (degree = 26), TRADD (degree = 26), RPS3 (degree = 26), AKT1 (degree = 25), YWHAB (degree = 24), HSPA1A (degree = 24), RPS3A (degree = 24) presenting the highest degree, suggesting that these proteins may play an essential role in maintaining the local connectivity of LRRK2_net_.

**Figure 4.**
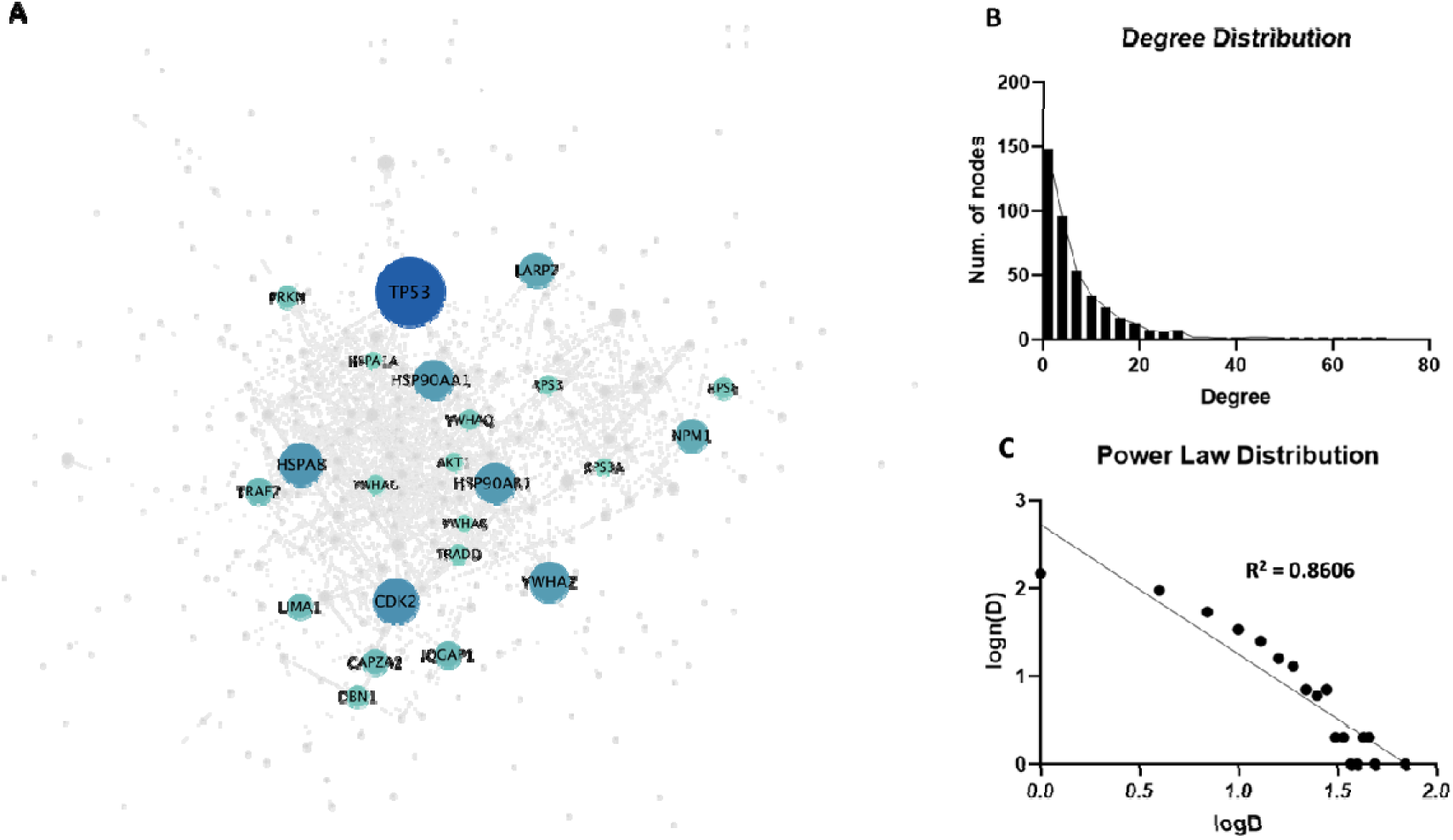
LRRK2_net_. **A)** The network graph shows the LRRK2 _net_, in which nodes represent the LRRK2 interactors (N = 338), while edges represent the “2-layer” PPIs (N = 1345). Node size refers to the node degree. Interactors with higher centrality (with degree ≥ 24) were colour-coded according to their degree. **B)** The bar graph shows the distribution of degree of LRRK2 interactors in the network; **C)** The log-log plot shows that the LRRK2_net_ follows the power law, in which the X-axis represents the log-transformed degree (logD), while the Y-axis represents the log-transformed frequency of a LRRK2 interactor with a certain degree level (log(n(D))). The scatters fit a linear regression line with R-square = 0.8606.

### Weighted network analysis on the LRRK2**_net_**

A total of 14 topological clusters were identified in the trimmed-LRRK2_net_ using the Fast Greedy algorithm based on the measure of edge betweenness (Figure 5A, **Table S7**). Of note, 3 clusters containing less than 5 interactors each were removed, leaving a total of 11 clusters for further analysis. For each of the 11 topological clusters, edges were classified as up/down-regulated or unchanged bases on the differential expression and co-expression levels of LRRK2 interactors in the sPD and LRRK2-PD conditions as compared to the controls. The distribution of the edges across these 3 categories was compared via One Sample Proportion Test to identified clusters significantly altered in expression in sPD or LRRK2-PD in comparison with controls (Figure 5B, **C**).

**Figure 5.**
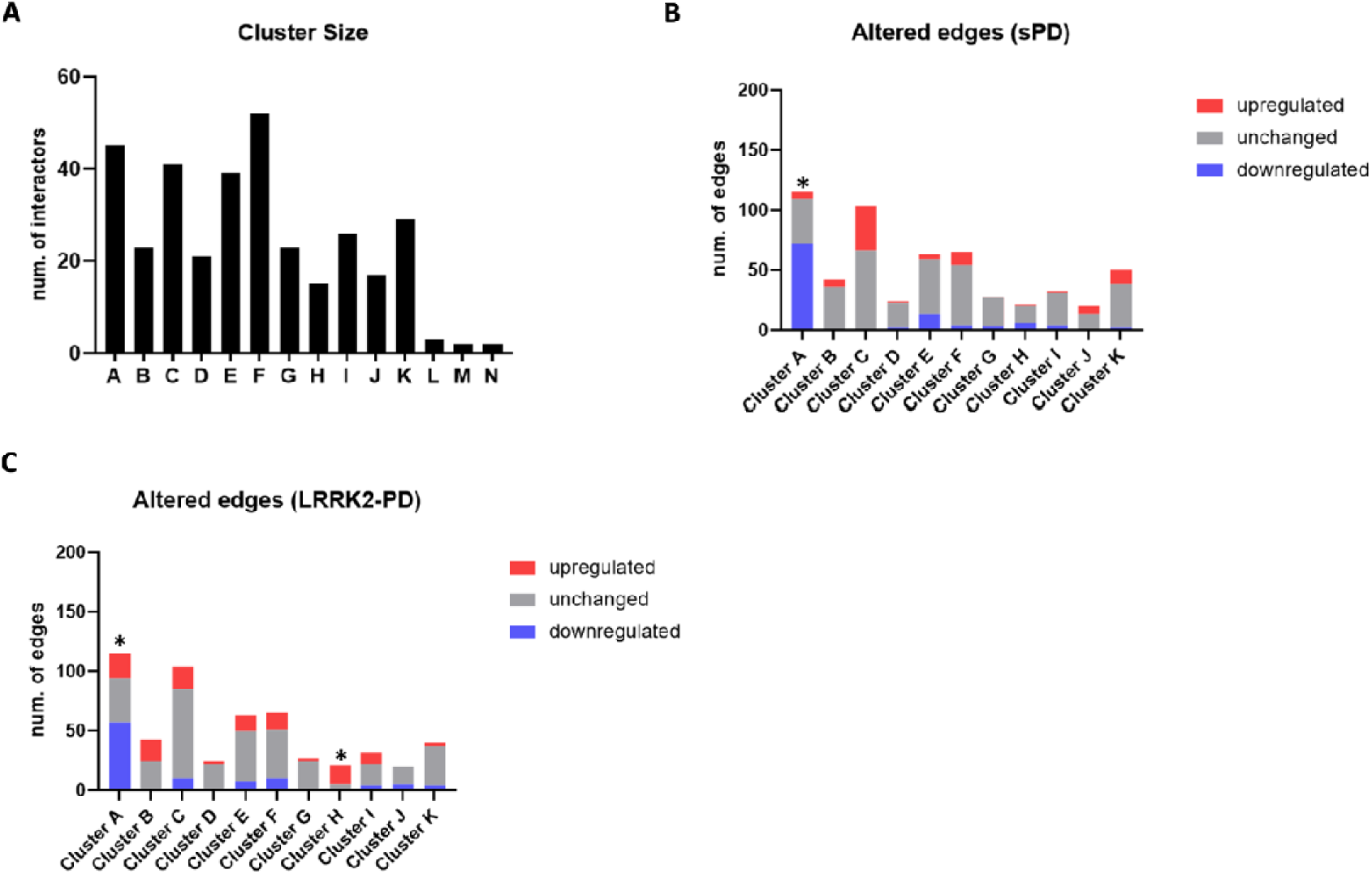
Topological clustering of the LRRK2 _net_. **A)** The bar graph shows the 14 topological clusters identified in the LRRK2net via the Fast Greedy Algorithm. Cluster L, M, N were discarded from further analysis due to their small size (contained ≤ 5 interactors). **B)** The bar graph shows the impact of expression behaviour linked with the sPD condition on the edges of each topological cluster. Upregulated edges (in red) were defined as: 1) with ≥ 1 connected interactor exhibiting increased expression level in sPD condition as compared to controls; and/or 2) the 2 connected interactors were positively co-expressed (with Pearson’ s coefficient > 0.6) in sPD condition but not in controls. Downregulated edges (in blue) were defined in the opposite way: 1) with ≥ 1 connected interactor exhibiting decreased expression level in sPD condition as compared to controls; and/or 2) the 2 connected interactors were positively co-expressed (with Pearson’s coefficient > 0.6) in controls but not in sPD. The percentage of upregulated, unchanged and downregulated edges were compared within each cluster via One Sample Proportion test. Only Cluster A was significantly downregulated in sPD (p < 0.001, *). **C)** The bar graph shows the impact of expression behaviour linked with the LRRK2-PD condition on the edges of each topological cluster. Cluster A was significantly downregulated in LRRK2-PD (p < 0.05, *); while Cluster H was significantly upregulated by LRRK2-PD, (p < 0.001, *) in comparison with controls.

Among the 11 clusters, Cluster A was significantly altered (down-regulated) in both sPD and LRRK2-PD cases vs. controls (p < 0.05), with 72/115 (62.6%) and 57/115 (49.5%) edges down-regulated, respectively (Figure 5B**,C**, Figure 6A). Functional enrichment analysis associated Cluster A with gene translation and ribosomal functions, suggesting that the sPD and LRRK2-PD pathologies potentially contribute to a perturbed ribosomal translation by down-regulating this cluster of LRRK2 interactors (Figure 6B, **Table S8**). In addition, Cluster H exhibited a significant up-regulation in the LRRK2-PD cases only, with a total of 13/21 (76.2%) edges up-regulated (Figure 5C, Figure 7A). Of note, up-regulated PRKN expression level contributed to 10/16 altered edges due to its central role in the cluster (degree = 11). Functional enrichment analysis related Cluster H to mitophagy and neurotransmitter transport, suggesting that during LRRK2-PD progression, regulation of these biological processes is potentially stimulated. (Figure 7B, **Table S8**).

**Figure 6.**
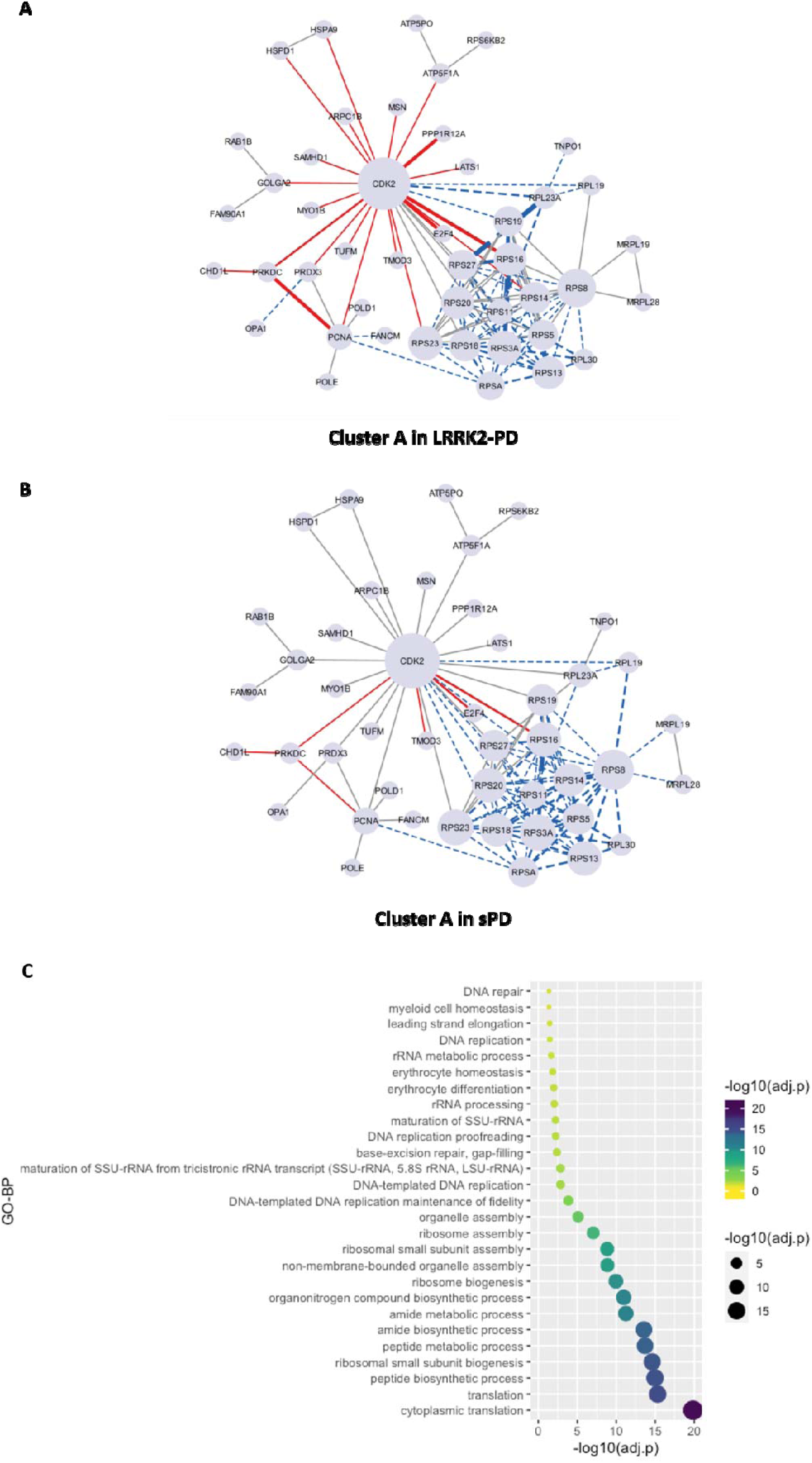
Significantly altered Topological Cluster A of the LRRK2 _net_ in the LRRK2-PD and sPD cases. **A)** The network graph shows the significant downregulation of Cluster A in sPD and LRRK2-PD, in which LRRK2 interactors are represented as nodes (N = 45) while PPIs are represented as edges (N = 115). Edges are represented with a continuous red line if they are up-regulated, with a dotted blue line if they are down-regulated. The thickness of the edges refers to the level of alterations of PPIs, i.e., if 2 interactors connected by a given edge exhibited the same trend of alteration (i.e, both up-regulated or down-regulated in PD cases vs. controls), the edge was marked thicker. **B)** The bubble graph shows the GO-BP terms enriched for the 45 LRRK2 interactors in Cluster A. Bubble colour and size refer to the enrichment significance (evaluated as -log10 transformed adjusted p-values, -log10(adj.p)).

**Figure 7.**
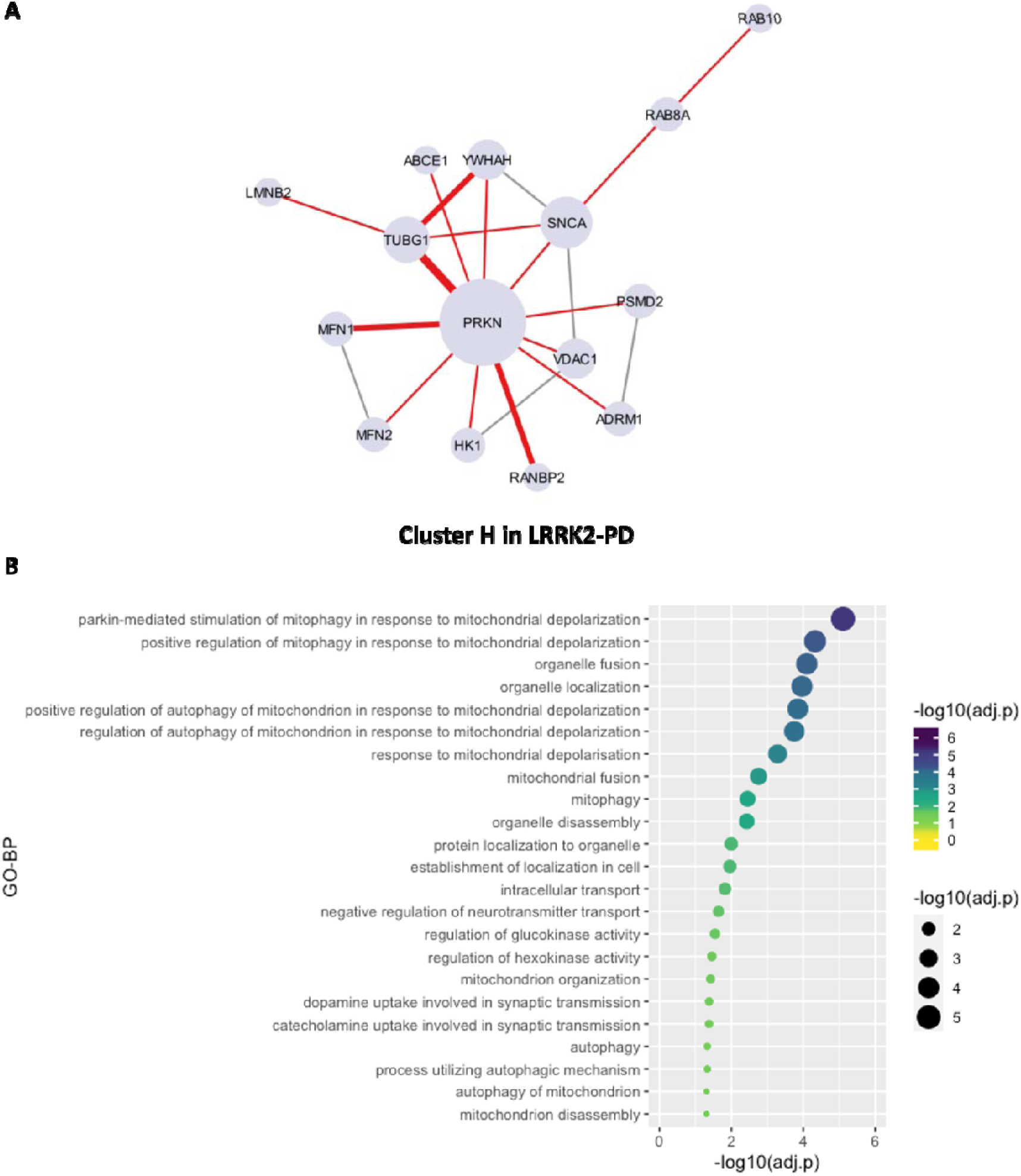
Significantly altered Topological Cluster H of the LRRK2 _net_ in the LRRK2-PD cases. **A)** The network graph shows the significant upregulated Cluster H in LRRK2-PD, with interactors represented as nodes (N = 15) and PPIs represented as edges (N = 21). Edges are represented with a continuous red line if they are up-regulated, with a dotted blue line if they are down-regulated. The thickness of the edges refers to the level of alterations of PPIs, i.e., if 2 interactors connected by a given edge exhibited the same trend of alteration (i.e, both up-regulated or down-regulated in PD cases vs. controls), the edge was marked thicker. **B)** The bubble graph shows the GO-BP terms enriched for the 15 LRRK2 interactors in Cluster H. Bubble colour and size refer to the enrichment significance (evaluated as -log10 transformed adjusted p-values, -log10(adj.p)).

## Discussion

Complex neurodegenerative disorders present with a complicated aetiopathogenesis, triggered by multiple causative events (or risk factors) from the environment and from the genome. An additional layer of complexity is due to the cocktail of risk factors being cohort specific. In PD the majority of cases present with a sporadic form of the disease, with no large effect size mutations contributing to it; these cases are considered to be due to a complex set of small effect size genetic risk factors in combination with a triggering environmental exposure. In contrast, a minority of patients is considered familial with at least one mutation with an effect size large enough to drive disease. The sporadic and the genetic forms of PD are, therefore, likely to be triggered by different combinations of risk factors. It is legitimate to ask whether, despite the similar clinical presentation and the classification under the same disease name, sporadic and genetic forms might represent a more nuanced spectrum of disorders. This question holds the key to a very practical issue. Sporadic disorders are difficult to model *in vitro*, thus the scientific community relies on genetic models based on the familial forms of the same disease to simulate the disease scenario in a test tube. These experimental models might not be accurate if we are indeed dealing with a spectrum disorder where the same clinical manifestation may be triggered by different genetic architectures and/or molecular scenarios. Similarly, a therapeutic approach targeted to the molecular core of the neurodegeneration developed for the genetic forms of the disease might not be fully effective on the sporadic disease, thus requiring cohort specific interventions.

In this study, we used a systems biology approach to generate a model and investigate the potential molecular differences between sPD and LRRK2-PD, focusing on the transcriptomic expression profile of the LRRK2 protein interactome. It is known that LRRK2 interaction behaviour is affected by the presence of mutations (Manzoni et al., 2015); we therefore hypothesised that the presence of PD causing mutations in LRRK2 (LRRK2-PD) will modify the LRRK2 connectivity and in turn this will trigger expression changes within the LRRK2 interactome. These changes might be specific of the LRRK2-PD scenario since no LRRK2 mutations are present in sPD. However, it is also possible that expression changes of the LRRK2 interactome might happen just as a consequence of PD, in a feedback response to the molecular alterations induced by the disease; in this case these alterations should be evident in both presence (LRRK2-PD) and absence (sPD) of LRRK2 mutations.

When we checked for alterations in expression levels (cases vs controls) for the LRRK2 interactors, 26.5% (100/377) of the LRRK2 interactors presented indeed significant alteration in whole blood mRNA, among which only 17 (5.6%) showed similar trend of alteration (3 up-regulated and 14 down-regulated) between sPD and LRRK2-PD. Functional enrichment of these 17 proteins with similar trend of alteration in both sPD and LRRK2-PD indicated biological processes related to ribosomal activity and protein biosynthesis. It is interesting to note that, among the 100 LRRK2 interactors that were significantly altered in expression levels (all cases vs controls), 23 were ribosomal proteins (RP). These RPs were all down-regulated in the disease scenario: 6 were down-regulated in LRRK2-PD only; 8 were down-regulated in sPD only while 9 were down-regulated in both sPD and LRRK2-PD. LASSO regression modelling showed that the expression features of these interactors were able to differentiate the LRRK2-PD and sPD cohort with a validated accuracy of 68%. This first analysis of the model suggested that, in general, sPD and LRRK2-PD possess distinct molecular signature of expression changes for the LRRK2 interactors. However, a small overlap in expression change does exist between the 2 conditions and this seems associated with those LRRK2 interactors that aid LRRK2 in its functions in the regulation of protein synthesis and ribosomal activity.

We then analysed the co-expression behaviour of the LRRK2 interactome using the WGCNA pipeline to identify modules of LRRK2 interactors with similar expression behaviour (co-expression) across the sPD, LRRK2-PD and control cohorts. A total of 2 co-expression modules were identified that consistently exist in the 3 conditions, which suggest functional units of LRRK2 interactors that participate in communal processes. Module-Trait analyses found that one of the 2 modules (the Turquoise module) was down-regulated in both sPD and LRRK2-PD cases as compared to controls, while the other one (the Blue module) remained unchanged in heath vs disease scenario. Interestingly, no co-expression module was identified containing LRRK2 itself, suggesting that in blood, at basal level, no LRRK2 interactor is significantly co-expressed with LRRK2.

We finally proceed to identify topological clusters within the LRRK2 interactome, based on the protein connections across LRRK2 interactors. Topological clusters might indicate functional local communities within a larger network based on how proteins relate/connect with each other. The topological clustering algorithm identified 11 clusters in the LRRK2_net_, these are portions of the network that are more connected within each other than the average connection of the entire network. The majority of the RP were, as expected, contained within one cluster and interestingly this cluster was the only one whose global change in expression and co-expression was overall statistically significant (cases vs controls) in both sPD and LRRK2-PD. In particular this cluster was significantly downregulated, potentially suggesting that the functionality of RPs and ribosomal protein biosynthetic processes are universally reduced during PD. This finding is in accordance with several previous functional studies in which similar down-regulation in RPs were found in whole blood and substantia nigra in sPD patients as well as LRRK2-G2019S PD patients as compared to controls (Scherzer *et al*., 2007; Garcia-Esparcia *et al*., 2015; Flinkman *et al*., 2023; Jang *et al*., 2023). The other network cluster that was significantly modified considering its expression behaviour in cases vs controls contained 15 LRRK2 interactors with general up-regulation; however, this cluster was only significantly modified in LRRK2-PD, but not in sPD when compared to control. The function of this cluster was associated with the mitochondria functions and mitophagy, and indeed PRKN/Parkin was at the centre of the cluster connectivity suggesting an interlink between of the effect of LRRK2 mutation (G2019S and/or R1441C) and the well-established mitophagy alterations in PD. Multiple lines of evidence have indicated that increased LRRK2 kinase activity negatively affects Parkin-dependent mitophagy. For example, a structural biology study has found that overexpression of LRRK2 and LRRK2-G2019S interrupts the PPI between Parkin and other key proteins essential for mitophagy on the outer mitochondrial membrane, and thereby interferes with the recruitment of Parkin with disruption of mitophagy (Bonello et al., 2019; Lin et al., 2021) This may explain the up-regulated PRKN/Parkin expression level cluster as observed in our study – as a compensation for a reduction in the available pool of Parkin free to interact at the mitochondria. In addition, the increased co-expression of PRKN/Parkin and MFN1, TUBG1 as well as RANBP2 may serve as compensation mechanism for the altered mitochondrial fusion/fission dynamics in the presence of LRRK2-G2019S, which have been reported in several studies (Bradshaw et al., 2021; Gegg et al., 2010). Additional functions associated with this LRRK2-PD only cluster were related to autophagy and, to a smaller extent, to neurotransmitter transport and synaptic transmission.

There are a number of limitations to this study: 1) the sample sizes of the cohorts are relatively small, especially for the LRRK2-PD cohort. Larger sample size would improve statistical power and thereby yield more robust results; 2) PD cases recruited by PPMI were at the early stages of the disease and the whole blood mRNA sequencing was run at the first visit. Therefore, the alterations of some LRRK2 interactors could be too subtle to be detected by DEA or WGCNA; 3) the LRRK2-PD cohort includes both G2019S and R1441C/G mutations and thereby introduce subtle bias due to genetic variation – inclusion of other coding variants associated with LRRK2-PD would strengthen the case for any changes being generalisable.

In conclusion, our study suggests that the molecular pathways at the basis of sPD and LRRK2-PD might be slightly different despite overlapping aspects of pathology and clinical presentation. There are shared changes of the LRRK2 interactome that can be appreciated at the transcriptome level in both the 2 conditions, with these predominantly associated with alterations of ribosomal proteins and proteins whose function is important for protein biosynthesis. However, there are also large differences between the 2 conditions suggested by the unique transcriptomics signatures. These findings suggest that LRRK2-PD and sPD should not be approached as the same conditions. This prompts the requirement for specific experimental models to be generated to differentially study sporadic and LRRK2 PD and confirms the requirement for careful patient stratification in clinical trials.

## Supplementary tables

**Table S1.** The LRRK2 interactome

**Table S2.** DEA of LRRK2 interactors in the sPD cases vs controls and the LRRK2-PD cases vs controls

**Table S3.** Functional enrichment results for the differentially expressed LRRK2 interactors (UP or DOWN-regulated) for LRRK2-PD vs controls and sPD vs controls

**Table S4.** Co-expression modules of LRRK2 interactors as identified by WGCNA

**Table S5.** Functional enrichment results for the co-expression modules identified by WGCNA

**Table S6.** 2^nd^-layer interactions across LRRK2’s direct interactors obtained from HIPPIE

**Table S7.** Clusters in the 2^nd^ layer LRRK2 protein interaction network

**Table S8.** Functional enrichment results for the Topological Cluster A and Cluster H in the LRRK2 protein interaction network

**Figure S1.**
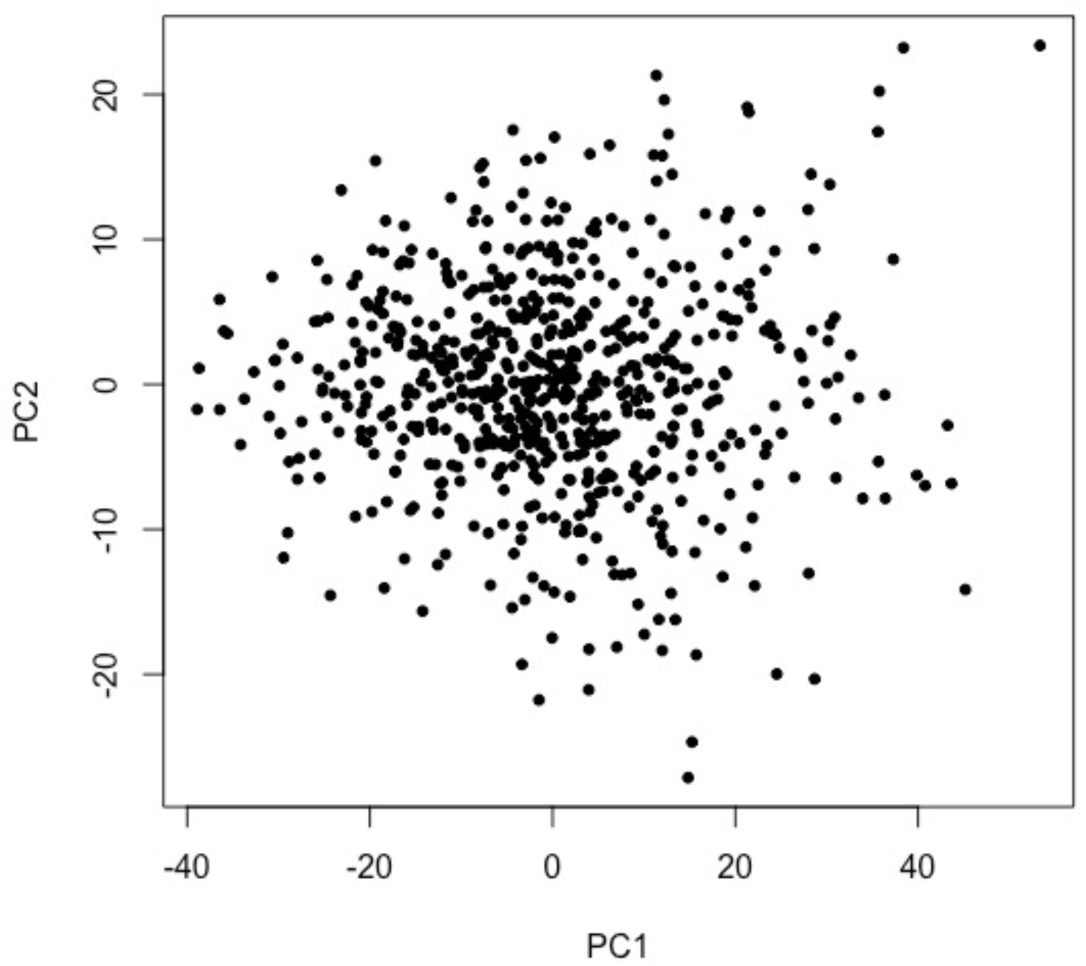
Subject QC on PPMI cohort. PCA was performed on the matrix of whole blood mRNA levels of the LRRK2 interactors for the 3 PPMI cohorts (control, sPD and LRRK2-PD). No outliers were excluded from further analysis

## Data Availability

Data used in the preparation of this article were obtained [on 24th January 2023] from the Parkinson’s Progression Markers Initiative (PPMI) database (www.ppmi-info.org/access-data-specimens/download-data), RRID:SCR_006431. For up-to-date information on the study, visit www.ppmi-info.org.

## Acknowledgments

This research was funded by the MJFF foundation (MJFF-023427 to CM and KH). CM and VEP receive funding from the MJFF (grant number MJFF-021335). PAL and CM acknowledge funding from the Biomarkers Across Neurodegenerative Diseases Grant Program 2019, BAND3 (Michael J. Fox Foundation, Alzheimer’s Association, Alzheimer’s Research UK, and the Weston Brain Institute [grant number 18063]). Funding: PPMI – a public-private partnership – is funded by the Michael J. Fox Foundation for Parkinson’s Research and funding partners, including 4D Pharma, Abbvie, AcureX, Allergan, Amathus Therapeutics, Aligning Science Across Parkinson’s, AskBio, Avid Radiopharmaceuticals, BIAL, Biogen, Biohaven, BioLegend, BlueRock Therapeutics, Bristol-Myers Squibb, Calico Labs, Celgene, Cerevel Therapeutics, Coave Therapeutics, DaCapo Brainscience, Denali, Edmond J. Safra

Foundation, Eli Lilly, Gain Therapeutics, GE HealthCare, Genentech, GSK, Golub Capital, Handl Therapeutics, Insitro, Janssen Neuroscience, Lundbeck, Merck, Meso Scale Discovery, Mission Therapeutics, Neurocrine Biosciences, Pfizer, Piramal, Prevail Therapeutics, Roche, Sanofi, Servier, Sun Pharma Advanced Research Company, Takeda, Teva, UCB, Vanqua Bio, Verily, Voyager Therapeutics, the Weston Family Foundation and Yumanity Therapeutics.

## Notes

### Competing Interest Statement

The authors have declared no competing interest.

### Summary of Updates

WGCNA analysis was modified from the previous version and we added a machine learning model via Least Absolute Shrinkage and Selection Operator (LASSO) algorithm

